# Discovery of the first genome-wide significant risk loci for ADHD

**DOI:** 10.1101/145581

**Authors:** Ditte Demontis, Raymond K. Walters, Joanna Martin, Manuel Mattheisen, Thomas D. Als, Esben Agerbo, Rich Belliveau, Jonas Bybjerg-Grauholm, Marie Bækvad-Hansen, Felecia Cerrato, Kimberly Chambert, Claire Churchhouse, Ashley Dumont, Nicholas Eriksson, Michael Gandal, Jacqueline Goldstein, Jakob Grove, Christine S. Hansen, Mads E. Hauberg, Mads V. Hollegaard, Daniel P. Howrigan, Hailiang Huang, Julian Maller, Alicia R. Martin, Jennifer Moran, Jonatan Pallesen, Duncan S. Palmer, Carsten B. Pedersen, Marianne G. Pedersen, Timothy Poterba, Jesper B. Poulsen, Stephan Ripke, Elise B. Robinson, Kyle F. Satterstrom, Christine Stevens, Patrick Turley, Hyejung Won, ADHD Working Group of the Psychiatric Genomics Consortium (PGC), Early Lifecourse & Genetic Epidemiology (EAGLE) Consortium, 23andMe Research Team, Ole A. Andreassen, Christie Burton, Dorret Boomsma, Bru Cormand, Søren Dalsgaard, Barbara Franke, Joel Gelernter, Daniel Geschwind, Hakon Hakonarson, Jan Haavik, Henry Kranzler, Jonna Kuntsi, Kate Langley, Klaus-Peter Lesch, Christel Middeldorp, Andreas Reif, Luis A. Rohde, Panos Roussos, Russell Schachar, Pamela Sklar, Edmund Sonuga-Barke, Patrick F. Sullivan, Anita Thapar, Joyce Tung, Irwin Waldman, Merete Nordentoft, David M. Hougaard, Thomas Werge, Ole Mors, Preben B. Mortensen, Mark J. Daly, Stephen V. Faraone, Anders D. Børglum, Benjamin M. Neale

## Abstract

Attention-Deficit/Hyperactivity Disorder (ADHD) is a highly heritable childhood behavioral disorder affecting 5% of school-age children and 2.5% of adults. Common genetic variants contribute substantially to ADHD susceptibility, but no individual variants have been robustly associated with ADHD. We report a genome-wide association meta-analysis of 20,183 ADHD cases and 35,191 controls that identifies variants surpassing genome-wide significance in 12 independent loci, revealing new and important information on the underlying biology of ADHD. Associations are enriched in evolutionarily constrained genomic regions and loss-of-function intolerant genes, as well as around brain-expressed regulatory marks. These findings, based on clinical interviews and/or medical records are supported by additional analyses of a self-reported ADHD sample and a study of quantitative measures of ADHD symptoms in the population. Meta-analyzing these data with our primary scan yielded a total of 16 genome-wide significant loci. The results support the hypothesis that clinical diagnosis of ADHD is an extreme expression of one or more continuous heritable traits.

## Background

Attention-Deficit/Hyperactivity Disorder (ADHD) is a neurodevelopmental psychiatric disorder, that affects around 5% of children and adolescents and 2.5% of adults worldwide^1^. ADHD is often persistent and markedly impairing with increased risk of harmful outcomes such as injuries^2^, traffic accidents^3^, increased health care utilization^4,5^, substance abuse^6^, criminality^7^, unemployment^8^, divorce^4^, suicide^9^, AIDS risk behaviors^8^, and premature mortality^10^. Epidemiologic and clinical studies implicate genetic and environmental risk factors that affect the structure and functional capacity of brain networks involved in behavior and cognition^1^, in the etiology of ADHD.

Consensus estimates from over 30 twin studies indicate that the heritability of ADHD is 70-80% throughout the lifespan^11,12^ and that environmental risks are those not shared by siblings^13^. Twin studies also suggest that diagnosed ADHD represents the extreme tail of one or more heritable quantitative traits^14^. Additionally, family and twin studies report genetic overlap between ADHD and other conditions including antisocial personality disorder/behaviours^15^, cognitive impairment^16^, autism spectrum disorder^17,18^, schizophrenia^19^, bipolar disorder^20^, and major depressive disorder^21^.

Thus far genome-wide association studies (GWASs) to identify common DNA variants that increase the risk of ADHD have not been successful^22^. Nevertheless, genome-wide SNP heritability estimates range from 0.10 – 0.28^23,24^ supporting the notion that common variants comprise a significant fraction of the risk underlying ADHD^25^ and that with increasing sample size, and thus increasing statistical power, genome-wide significant loci will emerge.

Previous studies have demonstrated that the common variant risk, also referred to as the single nucleotide polymorphism (SNP) heritability, of ADHD is also associated with depression^25^, conduct problems^26^, schizophrenia^27^, continuous measures of ADHD symptoms^28,29^ and other neurodevelopmental traits^29^ in the population. Genetic studies of quantitative ADHD symptom scores in children further support the idea that ADHD is the extreme of a quantitative trait^30^. Here we present a genome-wide meta-analysis identifying the first genome-wide significant loci for ADHD using a combined sample of 55,374 individuals from an international collaboration. We also strengthen the case that the clinical diagnosis of ADHD is the extreme expression of one or more heritable quantitative traits, at least as it pertains to common variant genetic risk, by integrating our results with previous GWAS of ADHD-related behavior in the general population.

## Genome-wide significantly associated ADHD risk loci

Genotype array data for 20,183 ADHD cases and 35,191 controls were collected from 12 cohorts (Supplementary Table 1). These samples included a population-based cohort of 14,584 cases and 22,492 controls from Denmark collected by the Lundbeck Foundation Initiative for Integrative Psychiatric Research (iPSYCH), and 11 European, North American and Chinese cohorts aggregated by the Psychiatric Genomics Consortium (PGC). ADHD cases in iPSYCH were identified from a national research register and diagnosed by psychiatrists at a psychiatric hospital according to ICD10 (F90.0), and genotyped using Illumina PsychChip. Designs for the PGC cohorts has been described previously^24,25,31,32,22^ (see Supplementary Information for detailed cohort descriptions).

Prior to analysis, stringent quality control procedures were performed on the genotyped markers and individuals in each cohort using a standardized pipeline^33^ (see Online Methods). Related individuals were removed, and genetic outliers were excluded based on principal component analysis. Non-genotyped markers were imputed using the 1000 Genomes Project Phase 3 reference panel^34^ (see Online Methods).

GWAS was conducted in each cohort using logistic regression with the imputed additive genotype dosages. Principal components were included as covariates to correct for population stratification^35^ (see Supplementary Information), and variants with imputation INFO score < 0.8 or minor allele frequency (MAF) < 0.01 were excluded. The GWAS were then meta-analyzed using an inverse-variance weighted fixed effects model^36^. Association results were considered only for variants with an effective sample size greater than 70% of the full meta-analysis, leaving 8,047,421 variants in the final meta-analysis. A meta-analysis restricted to European-ancestry individuals (19,099 cases, 34,194 controls) was also performed to facilitate secondary analyses.

In total, 304 genetic variants in 12 loci surpassed the threshold for genome-wide significance (P<5×10^−8^; Figure 1, Table 1, Supplementary Figure 3.A2 – 3.M2). No marker demonstrated significant heterogenety between studies (Supplementary Figure 6 and 7). Conditional analysis within each locus did not identify any independent secondary signals meeting genome-wide significance (see Online Methods, Supplementary Table 2).

**Figure 1.**
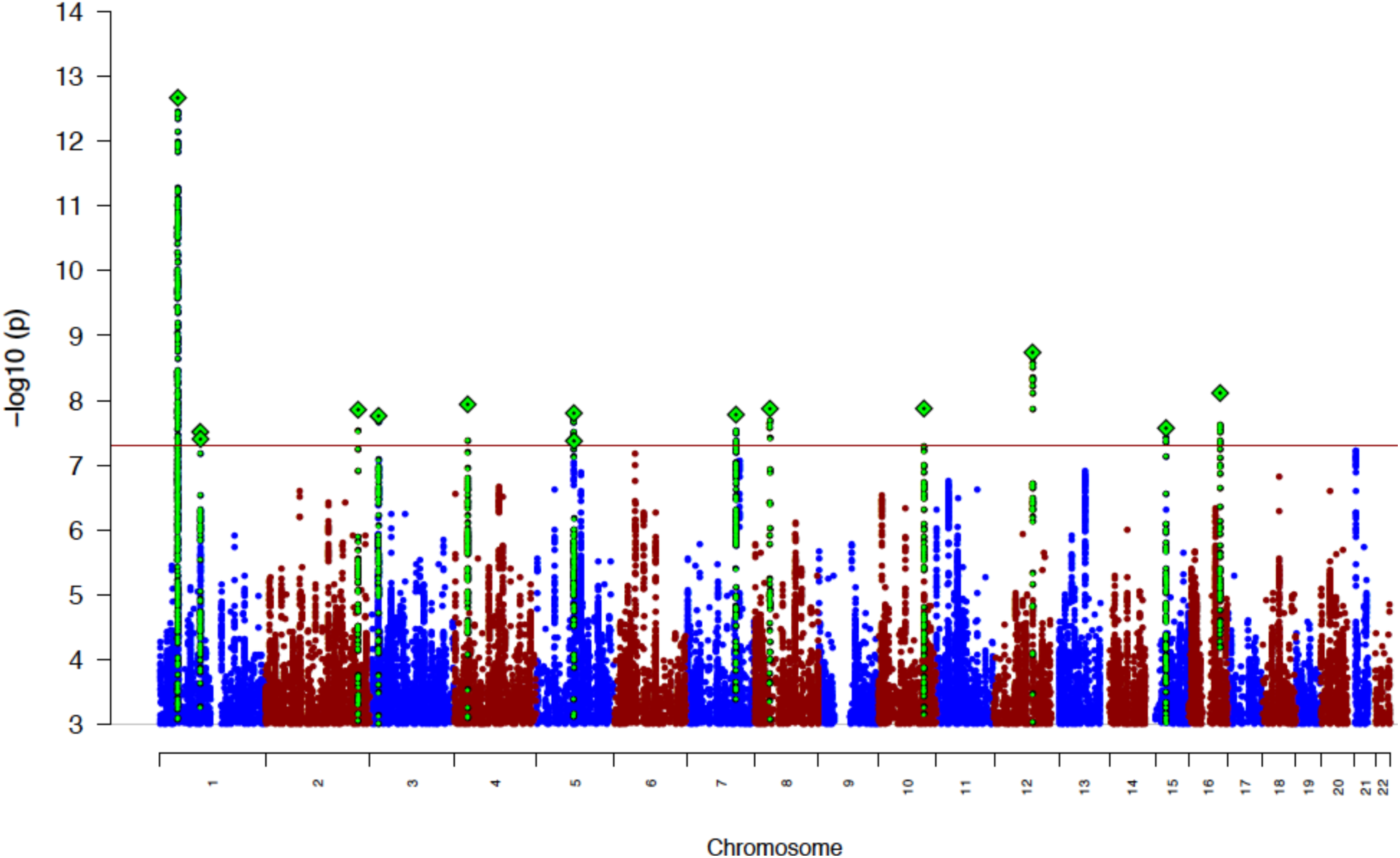
Manhattan plot of the results from the GWAS meta-analysis of ADHD. The index variants in the 12 genome-wide significant loci are highlighted as a green diamond. Index variants located with a distance less than 400kb are considered as one locus.

**Table 1.** Results for the genome-wide significant index variants in the 12 loci associated with ADHD identified in the GWAS meta-analysis. Index variants are LD independent (r^2^ < 0.1), and are merged into one locus when located with a distance less than 400kb. The location (chromosome [Chr] and base position [BP]), alleles (A1 and A2), allele frequency (A1 Freq), odds ratio (OR) of the effect with respect to A1, and association P-value of the index variant are given, along with genes within 50kb of the credible set for the locus.

| Locus | Chr | BP | Index Variant | Genes | A1 | A2 | A1 Freq | OR | P-value |
| --- | --- | --- | --- | --- | --- | --- | --- | --- | --- |
| 1 | 1 | 44184192 | rs11420276 | *ST3GAL3, KDM4A, KDM4A-AS1, PTPRF, SLC6A9, ARTN, DPH2, ATP6V0B, B4GALT2, CCDC24, IPO13* | G | GT | 0.696 | 1.113 | $2.14 \times 10^{-13}$ |
| 2 | 1 | 96602440 | rs1222063 | Intergenic | A | G | 0.328 | 1.101 | $3.07 \times 10^{-8}$ |
| 3 | 2 | 215181889 | rs9677504 | *SPAG16* | A | G | 0.109 | 1.124 | $1.39 \times 10^{-8}$ |
| 4 | 3 | 20669071 | rs4858241 | Intergenic | T | G | 0.622 | 1.082 | $1.74 \times 10^{-8}$ |
| 5 | 4 | 31151456 | rs28411770 | *PCDH7, LINC02497* | T | C | 0.651 | 1.09 | $1.15 \times 10^{-8}$ |
| 6 | 5 | 87854395 | rs4916723 | *LINC00461, MIR9-2, LINC02060, TMEM161B-AS1* | A | C | 0.573 | 0.926 | $1.58 \times 10^{-8}$ |
| 7 | 7 | 114086133 | rs5886709 | *FOXP2, MIR3666* | G | GTC | 0.463 | 1.079 | $1.66 \times 10^{-8}$ |
| 8 | 8 | 34352610 | rs74760947 | *LINC01288* | A | G | 0.957 | 0.835 | $1.35 \times 10^{-8}$ |
| 9 | 10 | 106747354 | rs11591402 | *SORCS3* | A | T | 0.224 | 0.911 | $1.34 \times 10^{-8}$ |
| 10 | 12 | 89760744 | rs1427829 | *DUSP6, POC1B* | A | G | 0.434 | 1.083 | $1.82 \times 10^{-9}$ |
| 11 | 15 | 47754018 | rs281324 | *SEMA6D* | T | C | 0.531 | 0.928 | $2.68 \times 10^{-8}$ |
| 12 | 16 | 72578131 | rs212178 | *LINC01572* | A | G | 0.883 | 0.891 | $7.68 \times 10^{-9}$ |

## Homogeneity of effects between cohorts

No genome-wide significant heterogeneity was observed in the ADHD GWAS meta-analysis (see Supplementary Information). Genetic correlation analysis (see Online Methods) provided further evidence that effects were consistent across different cohort study designs. The estimated genetic correlation between the European ancestry PGC samples and the iPSYCH sample from LD score regression^37^ was not significantly less than 1 (r_g_ = 1.17, SE = 0.2). The correlation between European ancestry PGC case/control and trio cohorts estimated with bivariate GREML was close to one (r_g_ = 1.02; SE = 0.32).

Polygenic risk scores (PRS)^38^ also show consistency over target samples. PRS computed in each PGC study using iPSYCH as the training sample were consistently higher in ADHD cases as compared to controls or pseudo-controls (see Supplementary Figure 11). Increasing deciles of PRS in the PGC were associated with higher odds ratio (OR) for ADHD (Figure 2). A similar pattern was seen in five-fold cross validation in the iPSYCH sample, with PRS for each subset computed from the other four iPSYCH subsets and the PGC samples used as training samples (see Online Methods; Figure 2). Across iPSYCH subsets, the mean of the maximum variance explained by the estimated PRS (Nagelkerke’s R^2^) was 5.5% (SE = 0.0012). The difference in standardized PRS between cases and controls was stable across iPSYCH subsets (OR = 1.56, 95% confidence interval (CI): 1.53 – 1.60); Supplementary Figure 9). These results further support the highly polygenic architecture of ADHD and demonstrate that the risk is significantly associated with the individual PRS burden in a dose-dependent manner.

**Figure 2.**
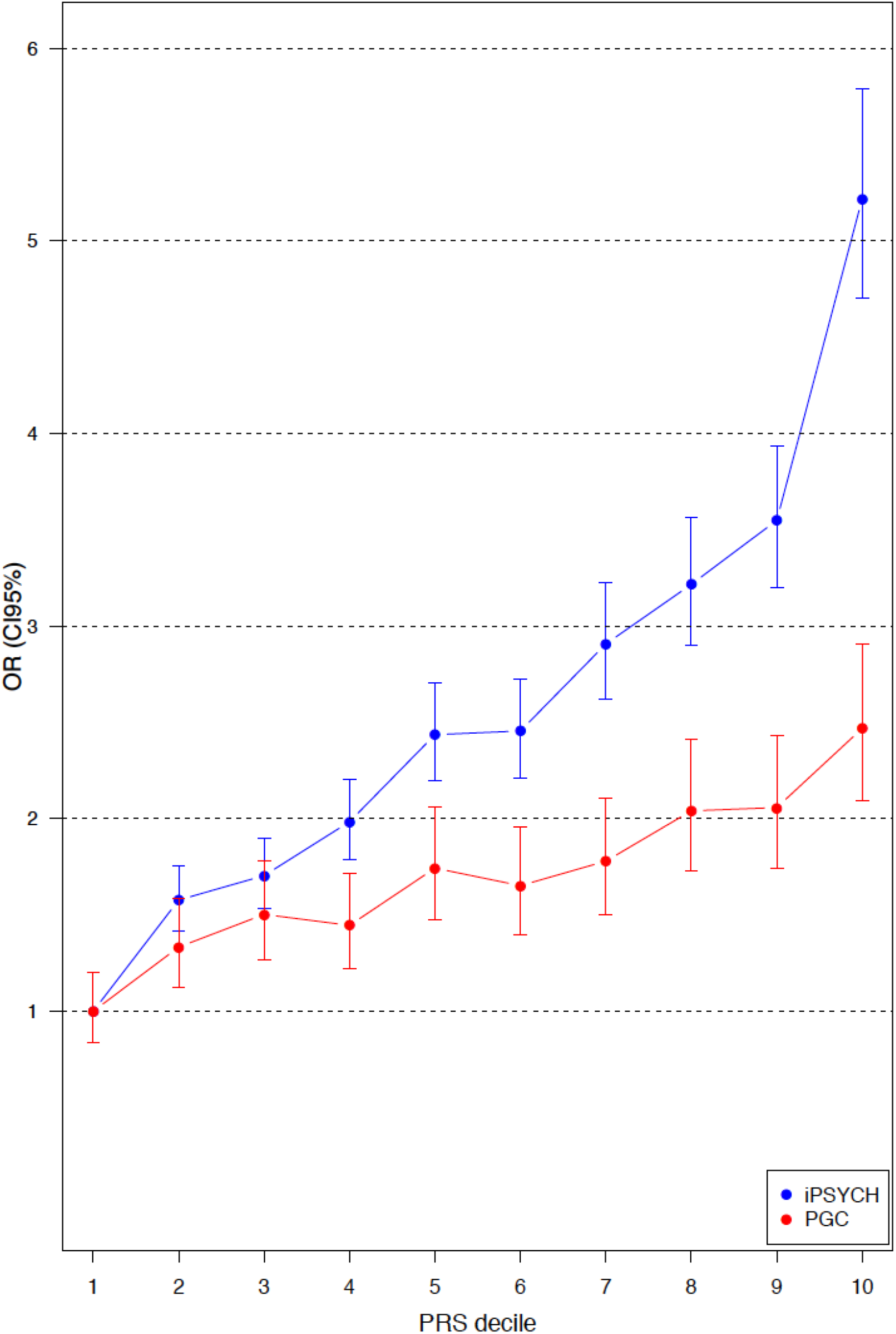
Odds Ratio (OR) by PRS within each decile estimated for individuals in the PGC samples (red dots) and in the iPSYCH sample (blue dots). Error bars indicate 95% confidence limits.

## Polygenic Architecture of ADHD

To assess the proportion of phenotypic variance explained by common variants we applied LD score regression^37^ in the European ancestry meta-analysis (Online Methods). Assuming a population prevalence of 5% for ADHD^39^, we estimate that the liability-scale SNP heritability h^2^_snp_=0.216 (SE=0.014; P=8.18×10^−54^). These estimated polygenic effects account for 88% (SE=0.0335) of observed genome-wide inflation of the test statistics in the meta-analysis; the remaining inflation, which may reflect confounding factors such as cryptic relatedness and population stratification, is significant but modest (intercept=1.0362, SE=0.0099; P=2.27×10^−4^).

To further characterize the patterns of heritability from the genome-wide association data, we performed partitioning based on the functional annotations described in Finucane et al.^40^ (see Online Methods). The analysis revealed significant enrichment in the heritability by SNPs located in conserved regions (P = 8.49 x 10^−10^) (Supplementary Figure 12), supporting the general biological importance of conserved regions and the potential impact on ADHD of variants located in these regions. Additionally enrichment in the heritability of SNPs located in cell-type-specific regulatory elements was evaluated by using the cell-type-specific group annotations described in Finucane et al^40^. The analysis revealed a significant enrichment of the average per SNP heritability for variants located in central nervous system specific regulatory elements (enrichment = 2.44 (SE=0.35); P = 5.81 x 10^−5^) (Supplementary Figures 13 and 14).

## Genetic correlation with other traits

Pairwise genetic correlation with ADHD was estimated for 220 phenotypes using LD score regression^41,42^ (Online Methods, Supplementary eTable 5). Thirty-eight phenotypes demonstrated significant genetic overlap with ADHD (P<2.27×10^−4^), including major depressive disorder(submitted), educational outcomes^43–46^, obesity-related phenotypes^47–52^, smoking^53–55^, reproductive success^56^, and mortality^57^ (Figure 3; Supplementary Table 11). In each case the genetic correlation is supported in GWAS of multiple related phenotypes. For the positive genetic correlation with major depressive disorder (r_g_=0.42, P= 7.38×10^−38^), we also observe a positive correlation with depressive symptoms (r_g_=0.45, P= 7.00×10^−19^), neuroticism (r_g_=0.26, P= 1.02x10^−8^) and a negative correlation with subjective well-being (r_g_=-0.28, P= 3.73x10^−9^). The positive genetic correlations with ever smoking (r_g_=0.48, P= 4.33x10^−16^) and with number of cigarettes smoked (r_g_=0.45, P= 1.07x10^−5^) are reinforced by significant positive correlation with lung cancer (r_g_=0.39, P= 6.35x10^−10^). Similarly, genetic correlations related to obesity include significant relationships with body mass index (BMI; r_g_= 0.26, P= 1.68x10^−15^), waist-to-hip ratio (r_g_=0.30, P= 1.16x10^−17^), childhood obesity (r_g_= 0.22, P= 3.29x10^−6^), HDL cholesterol (r_g_= −0.22, P= 2.44x10^−7^), and Type 2 Diabetes (r_g_= 0.18, P= 7.80x10^−5^). Additionally the negative correlation with years of schooling (rg = −0.53, P= 6.02x10^−80^) is supported by a negative genetic correlation with childhood IQ (−0.41, P= 1.24x10^−6^).

**Figure 3.**
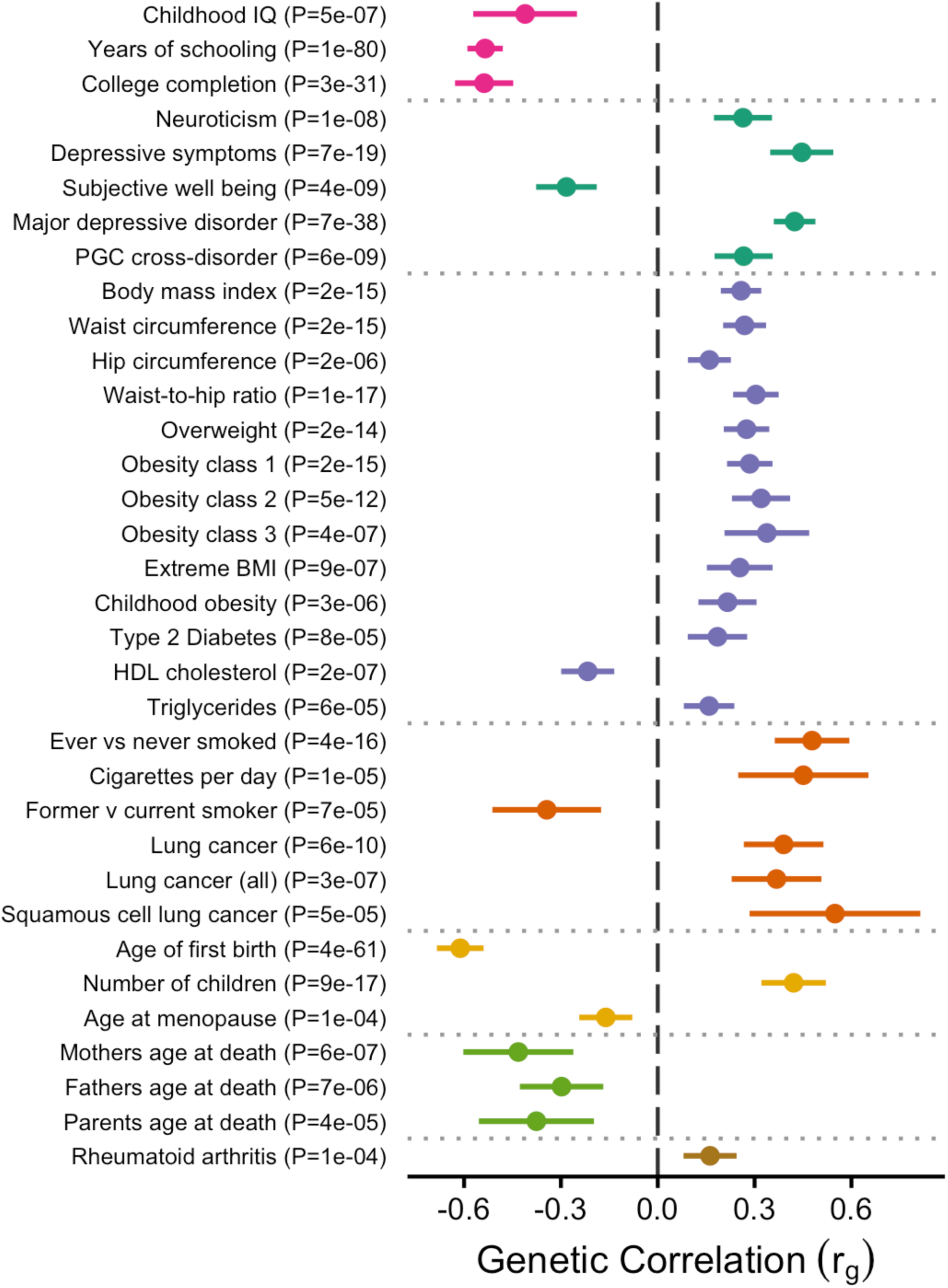
Significant genetic correlations between ADHD and other traits reveal overlap of genetic risk factors for ADHD across several groups of traits (grouping indicated by a horizontal line): educational, psychiatric/personality, weight (and possible weight related traits), smoking behaviour/smoking-related cancer, reproductive traits and parental longevity. In total 220 traits were tested. Two significant educational phenotypes are omitted due to substantial overlap with years of schooling. Error bars indicate 95% confidence limits.

## Biological annotation of significant loci

For the 12 genome-wide significant loci, Bayesian credible sets were defined to identify the set of variants at each locus most likely to include a causal effect (see Online Methods, Supplementary eTable 1). Biological annotations of the variants in the credible set were then considered to identify functional or regulatory variants, common chromatin marks, and variants associated with gene expression (eQTLs) or in regions with gene interactions observed in Hi-C data (see Online Methods, Supplementary eTable 2). Broadly, the significant loci do not coincide with candidate genes proposed to play a role in ADHD^58^.

Here we highlight genes that are identified in the regions of association (see also Supplementary Table 4). The loci on chromosomes 2, 7, and 10 each have credible sets localized to a single gene with limited additional annotations. In the chromosome 7 locus, *FOXP2* encodes a forkhead/winged-helix transcription factor and is known to play an important role in synapse formation and neural mechanisms mediating the development of speech and learning^59–61^. Comorbidity of ADHD with specific developmental disorders of language and learning is common (7-11%)^62,63^, and poor language skills have been associated with higher inattention/hyperactivity symptoms in primary school^64^.

Genome-wide significant loci on chromosomes 12 and 15 have biological annotations supporting the co-localized genes. The credible set on chromosome 12 spans *DUSP6*, and includes an annotated missense variant in the first exon and an insertion near the transcription start site, though neither is the lead variant in the locus (Supplementary eTable 3). The chromosome 15 locus is located in *SEMA6D*, and the majority of variants in the credible set are strongly associated with expression of *SEMA6D* in fibroblasts^65^. *DUSP6* encodes a dual specificity phosphatase^66^, and may play a role in regulating neurotransmitter homeostasis by affecting dopamine levels in the synapses^67,68^. Regulation of dopamine levels is likely to be relevant to ADHD since widely used ADHD medications have dopaminergic targets^69,70^ that increases the availability of synaptic dopamine. *SEMA6D* is active in the brain during embryonic development, and may play a role in neuronal wiring^71^. Variants in *SEMA6D* have previously been associated with eduational attainment^72^.

Credible set annotations at the remaining loci are more diverse (Supplementary eTable 2). The most strongly associated locus on chromosome 1 (index variant rs112984125) covers a gene-rich 250kb region of strong LD. The index variant is intronic to *ST3GAL3*, and most SNPs in the credible set are strongly associated with expression of *ST3GAL3* in whole blood^73^ (Supplementary eTable 2). Missense mutations in *ST3GAL3* have been shown to cause autosomal recessive intellectual disability^74^. Hi-C and eQTL annotations suggest multiple alternative genes however (Supplementary eTable 3). The locus also includes an intergenic variant, rs11210892, that has previously been associated with schizophrenia^33^.

On chromosome 5, the credible set includes links to *LINC00461* and *TMEM161B* (Supplementary eTable 2). The function of *LINC00461* is unclear, but the RNA has highly localized expression in the brain^75^ and the genome-wide significant locus overlaps with variants in *LINC00461* associated with educational attainment^72^. Alternatively, a genome-wide significant SNP in this locus (rs304132) is located in *MEF2C-AS1*, of strong interest given previous associations between *MEF2C* and severe intellectual disability,^76–78^ cerebral malformation^77^, depression^79^, schizophrenia^33^ and Alzheimer’s disease^80^, but the corresponding variant is not supported by the credible set analysis. Credible set annotations for other significant loci are similarly cryptic.

## Analysis of gene sets

Competitive gene based tests were performed for *FOXP2* target genes, highly constrained genes, and for all Gene Ontology terms^81^ from MsigDB 6.0^82^ using MAGMA^83^ (Online Methods). Association results for individual genes are consistent with the genome-wide significant loci for the GWAS (Supplementary Table 5). Three independent sets of *FOXP2* downstream target genes^84,85^ were tested (Online Methods), none of which demonstrated significant association to ADHD (Supplementary Table 7). The lack of association, might be caused by unknown functions of *FOXP2* driving ADHD risk; insufficient power to detect relevant downstream genes or that only a small subset of biological functions regulated by FOXP2 are relevant to ADHD pathogenesis.

Consistent with the partitioning of heritability, a set of 2,932 genes that are highly constrained and show high intolerance to loss of function^86^ showed significant association with ADHD (beta=0.062, P=2.6x10^−4^). We also find little evidence for effects in previously proposed candidate genes for ADHD^58^; of the nine proposed genes only *SLC9A9* showed weak association with ADHD (P=3.4x10^−4^; Supplementary Table 6). None of the Gene Ontology gene sets were significant after correction for multiple testing, but the top pathways did include interesting and nominally significant pathways such as “dopamine receptor binding” (p=0.0010) and “Excitatory Synapse” (P = 0.0088).(Supplementary eTable 4).

## Replication of GWAS loci

Here we describe the comparison of the GWAS meta-analysis of ADHD with two other ADHD-related GWASs: a 23andMe self-report cohort (5,857 cases and 70,393 controls) and a metaanalysis of childhood rating scales of ADHD symptoms performed by the EAGLE consortium (17,666 children < 13 years of age)^30^. We observed moderate concordance of genome-wide results between the ADHD GWAS and the cohort with a self-reported history of diagnosis for ADHD or Attention Deficit Disorder genotyped by 23andMe (see Supplementary Information). The estimated genetic correlation between the two analyses was strong (r_g_=0.653, SE=0.114), but significantly less than 1 (P=2.35x10^−3^). Of 94 clumped loci with P<1x10^−5^ in the ADHD GWAS meta-analysis, 71 had effects in the same direction in the 23andMe GWAS, significantly more concordant than expected by chance (P=1.25x10^−6^; Online Methods; Supplementary Table 12). We also observe a striking genetic correlation between the ADHD GWAS meta-analysis and quantitative measures of ADHD-related behavior in the general population as assembled by the EAGLE Consortium (r_g_=0.943, SE = 0.204). The direction of effect in EAGLE was concordant for 65 of 94 clumped loci with P<1x10^−5^ in the ADHD GWAS (P=3.06×10^−4^; Supplementary Table 12). For the 12 genome-wide significant loci in the ADHD GWAS meta-analysis, seven of the 11 loci present in the 23andMe GWAS had effects in the same direction (P=0.386 for sign concordance; Supplementary Table 13). In EAGLE, the direction of effect is concordant for 10 of 11 genome-wide significant loci from the ADHD GWAS meta-analysis (P =0.0159).

We then meta-analyzed the ADHD GWAS with the 23andMe and EAGLE results. For EAGLE, we developed a model to meta-analyze the GWAS of the continuous measure of ADHD with the clinical diagnosis in the ADHD GWAS. In brief, we perform a Z-score based meta-analysis using a weighting scheme based on the heritability and effective sample size for each phenotype (detailed description in Supplementary Information).

All 12 genome-wide significant loci from the ADHD GWAS meta-analysis maintained genome-wide significance after meta-analysis with EAGLE (Supplementary Figure 18), though the EAGLE data does not contribute to significance at two of those loci. Meta-analysis with EAGLE also yielded three additional loci surpassing genome-wide signficance (Supplementary eTable 6). Eight of the 12 ADHD loci were significant after inverse variance-weighted meta-analysis of the ADHD GWAS with 23andMe (Supplementary Figure 15), including two loci without 23andMe results. Joint meta-analysis of the ADHD, 23andMe, and EAGLE GWASs yielded significance for 10 of the 12 genome-wide significant ADHD loci and a total of 16 significant loci (Supplementary eTable 6, Supplementary Figure 21).

Consistent with the genetic correlation results, there is evidence of heterogeneity between the ADHD GWAS meta-analysis results and 23andMe GWAS. Genome-wide significant heterogeneity was observed at the lead chromosome 1 locus from the ADHD GWAS meta-analysis (rs12410155: I^2^=97.2, P=2.29×10^−9^; Supplementary Figure 17). No genome-wide significant heterogeneity was observed in the meta-analysis of ADHD and EAGLE (Supplementary Figure 20).

## Discussion

GWAS meta-analysis of ADHD revealed the first genome-wide significant risk loci, and indicates an important role for common variants in the polygenic architecture of ADHD. Several of the loci are located in or near genes that implicate neurodevelopmental processes that are likely to be relevant to ADHD, including *FOXP2* and *DUSP6*. Future work may focus on refining the source of the strong association in each locus, especially the lead locus on chromosome 1 which is complicated by broad LD and substantial heterogeneity between ADHD meta-analysis and analysis of self-reported ADHD status in 23andMe.

The 12 significant loci are compelling, but only capture a tiny fraction of common variant risk for ADHD. The odds ratios for the risk increasing allele at the index SNPs in the 12 significant loci are modest, ranging from 1.077 to 1.198 (Table 1). This is within the range of effect sizes for common genetic variants that has been observed for other highly polygenic psychiatric disorders e.g. schizophrenia^33^. A considerably larger proportion of the heritability of ADHD can be explained by all common variants (h^2^_snp_ = 0.22, SE = 0.01). This is consistent with previous estimates of h^2^_snp_ for ADHD in smaller studies (h^2^_snp_: 0.1-0.28)^23,24^, and also comparable to what has been found for schizophrenia (h^2^_snp_ 0.23 – 0.26)^23,24^. As would be hypothesized for a psychiatric disorder, these effects are enriched in conserved regions and regions containing enhancers and promoters of expression in the central nervous system tissues, consistent with previous observations in schizophrenia and bipolar disorder^40^.

Along with polygenicity, selection and evolutionary pressures may be an important feature of the architecture of ADHD genetics. We observe that ADHD risk variants are strongly enriched in genomic regions conserved in mammals^87^, and constrained genes likely to be intolerant of loss-of-function mutations^86^ are associated with ADHD. The directionality of potential selective effect is unclear: we find that common variant risk for ADHD is genetically correlated with having children younger and having more children, but is also correlated with a family history of parental mortality at a younger age. Given the documented association between ADHD and educational underachievement^88,89^, reinforced by genetic correlation of ADHD with educational attainment and childhood IQ^43^ observed in this study, selective pressure on the genetics of ADHD would be consistent with recent work suggesting that variants associated with educational attainment are under negative selection in Iceland^90^. Future studies of rare and *de novo* variants may provide more insight on selective pressures in ADHD-associated loci.

The observed genetic correlations with educational outcomes and other phenotypes suggest a strong genetic component to the epidemiological correlates of ADHD. The significant positive genetic correlation of ADHD with major depressive disorder and depressive symptoms supports previous findings suggesting a positive genetic overlap between those phenotypes^24,42^, as well as the broader genetic overlap of psychiatric disorders^23,24^. Positive genetic correlations between ADHD and health risk behaviors such as smoking and obesity are consistent with the observed increase in those behaviors among individuals with ADHD^91–94^ and indicates a shared genetic basis across these traits.

The current results further support the hypothesis that ADHD is the extreme expression of one or more heritable quantitative traits. We observe strong concordance between the GWAS of ADHD and previous GWAS of ADHD-related traits in the population from the EAGLE Consortium^30^, both in terms of genome-wide genetic correlation and concordance at individual loci. Polygenic risk for ADHD has previously been associated with inattentive and hyperactive/impulsive trait variation below clinical thresholds in the population^29^. Shared genetic risk with health risk behaviors may similarly be hypothesized to reflect an impaired ability to self-regulate and inhibit impulsive behavior^95,96^.

In summary, we report 12 independent genome-wide significant loci associated with ADHD in GWAS meta-analysis of 55,374 individuals from 12 study cohorts. The GWAS meta-analysis implicates *FOXP2*, *DUSP6*, and other constrained regions of the genome as important contributors to the eitiology of ADHD. The results also highlight strong overlap with the genetics of ADHD-related traits and health risk behaviors in the population, encouraging a dimensional view of ADHD.

## Online Methods

### GWAS meta-analysis

Quality control, imputation and primary association analyses were done using the bioinformatics pipeline Ricopili (available at https://github.com/Nealelab/ricopili), developed by the Psychiatric Genomics Consortium (PGC)^33^. In order to avoid potential study effects the 11 PGC samples and the 23 genotyping batches within iPSYCH were each processed separately unless otherwise stated (see Supplementary Information).

Stringent quality control was applied to each cohort following standard procedures for GWAS, including filters for call rate, Hardy-Weinberg equilibrium, and heterozygosity rates (see Supplementary Information). Each cohort was then phased and imputed using the 1000 Genomes Project phase 3 (1KGP3)^34,97^ imputation reference panel using SHAPEIT^98^ and IMPUTE2^99^, respectively. For trio cohorts, pseudocontrols were defined from phased haplotypes prior to imputation.

Cryptic relatedness and population structure were evaluated using a set of high quality markers pruned for linkage disequilibrium (LD). Genetic relatedness was estimated using PLINK v1.9^100,101^ to identify first and second-degree relatives 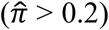 and one individual was excluded from each related pair. Genetic outliers were identified for exclusion based on principal component analyses using EIGENSOFT^35,102^. This was done separately for each of the PGC samples and on a merged set of genotypes for the iPSYCH sample (see Supplementary Information). Across studies, a total of 20,183 cases and 35,191 controls remained for analysis after QC.

Genome-wide association analyses for the 11 PGC samples and the 23 waves in iPSYCH were performed using logistic regression model with the imputed marker dosages in PLINK v1.9^100,101^. Principal components were included as covariates to control for population stratification^35,102^, along with relevant study-specific covariates where applicable (see Supplementary Information). Subsequently the results were meta-analysed using an inverse-variance weighted fixed effects model, implemented in METAL (version 2011-03-25) ^36^. Variants were filtered and included for imputation quality (INFO score) > 0.8 and MAF > 0.01. Only markers supported by an effective sample size N_eff_ = 2/(1/N_cases_ + 1/N_controls_)^103^ greater than 70% were included. After filtering, the meta-analysis included results for 8,047,421 markers.

### Conditional analysis

Twelve independent genome-wide significant loci were identified by LD clumping and merging loci within 400 kb (see Supplementary Information). In two of these loci a second index variant persisted after LD clumping. The two putative secondary signals were evaluated by considering analysis conditional on the lead index variant in each locus. In each cohort, logistic regression was performed with the imputed genotype dosage for the lead index variant included as a covariate. All covariates from the primary GWAS (e.g. principle components) were also included. The conditional association results were then combined in an inverse-variance weighted meta-analysis.

### Genetic correlations between ADHD samples

Genetic correlation between the European-ancestry PGC and iPSYCH GWAS results was calculated using LD Score regression^37^. The regression was performed using pre-computed LD scores for HapMap3 SNPs calculated based on 378 European-ancestry individuals from the 1000 Genomes Project (available on https://github.com/bulik/ldsc). Only results for markers with an imputation INFO score > 0.90 were included in the analysis. In addition, a bivariate GREML analysis was conducted using GCTA^104^ in order to estimate the genetic correlation between PGC case/control and trio study designs.

### Polygenic Risk Scores for ADHD

The iPSYCH sample were split into five groups, and subsequently five leave-one-out association analyses were conducted, using four out of five groups and the PGC samples as training datasets^38^. PRS were estimated for each target sample using variants passing a range of association P-value thresholds in the training samples. PRS were calculated by multiplying the natural log of the odds ratio of each variant by the allele-dosage (imputation probability) and whole-genome polygenic risk scores were obtained by summing values over variants for each individual.

For each of the five groups of target samples, PRS were normalized and the significance of the case-control score difference was tested by standard logistic regression including principle components. For each target group and for each P-value threshold the proportion of variance explained (i.e. Nagelkerke’s *R^2^*) was estimated by comparing the regression with PRS to a reduced model with covariates only. The OR for ADHD within each PRS decile group was estimated based on the normalized score across groups (using the P-value threshold with the highest Nagelkerke’s *R^2^* within each target group) (Figure 3). OR was also estimated using logistic regression on the continuous scores for each target group separately and an OR based on all samples using the normalized PRS score across all groups (Supplementary Figure 9). Additionally PRS were evaluated in the PGC samples using the iPSYCH sample as training sample, following the approach described above (see Supplementary Information).

### SNP heritability and intercept evaluation

LD score regression^37^ was used to evaluated the relative contribution of polygenic effects and confounding factors, such as cryptic relatedness and population stratification, to deviation from the null in the genome-wide distribution of GWAS *χ*^2^ statistics. Analysis was performed using pre-computed LD scores from European-ancestry samples in the 1000 Genomes Project (available on https://github.com/bulik/ldsc) and summary statistics for the European-ancestry ADHD GWAS to ensure matching of population LD structure. The influence of confounding factors was tested by comparing the estimated intercept of the LD score regression to one, it’s expected value under the null hypothesis of no confounding from e.g. population stratification. The ratio between this deviation and the deviation of the mean *χ*^2^ from one (i.e. it’s expected value under the null hypothesis of no association) was used to estimate the proportion of inflation in *χ*^2^ attributable to confounding as opposed to true polygenic effects (ratio = (intercept-1)/(mean *χ*^2^-1)). SNP heritability was estimated based on the slope of the LD score regression, with heritability on the liability scale calculated assuming a 5% population prevalence of ADHD^39^.

### Partitioning of the heritability

SNP heritability was partitioned by functional category and tissue association using LD score regression^40^. Partitioning was performed for 53 overlapping functional categories, as well as 220 cell-type-specific annotations grouped into 10 cell-type groups, as described in Finucane et al. ^40^. For both sets of annotations we used previously computed LD scores and allele frequencies from European ancestry samples in the 1000 Genomes Project (available on https://data.broadinstitute.org/alkesgroup/LDSCORE/).

Additionally we expanded the cell-type specific heritability analysis by including an annotation based on information about H3K4Me1 imputed gapped peaks excluding the broad MHC-region (chr6:25-35MB), generated by the Roadmap Epigenomics Mapping Consortium^105,106^ (see Supplementary Information). The analyses were restricted to the European GWAS meta-analysis results to ensure matching of population LD structure. Results for each functional category were evaluated based on marginal enrichment, defined as the proportion of SNP heritability explained by SNPs in the annotation divided by the proportion of genome-wide SNPs in the annotation^40^. For each cell-type group and each H3K4Me1 cell-type annotations, the contribution to SNP heritability was tested conditional on the baseline model containing the 53 functional categories.

### Genetic correlations of ADHD with other traits

The genetic correlation of ADHD with other traits were evaluated using LD Score regression^42^. For a given pair of traits, LD score regession estimates the expected population correlation between the best possible linear SNP-based predictor for each trait, restricting to common SNPs. Such correlation of genetic risk may reflect a combination of colocalization, pleiotropy, shared biological mechanisms, and causal relationships between traits. Correlations were tested for 219 phenotypes with publically available GWAS summary statistics using LD Hub^41^ (see Supplementary Information). Correlation with Major Depressive Disorder was tested using GWAS results from an updated analysis of 130,664 cases and 330,470 controls from the Psychiatric Genomics Consortium (submitted). As in the previous LD score regression analyses, this estimation was based on summary statistics from the European GWAS meta-analysis, and significant correlations reported are for traits analysed using individuals with European ancestry.

### Credible set analysis

We defined a credible set of variants in each locus using the method described by Maller et al.^107^ (see Supplementary Information), implemented by a freely available R script (https://github.com/hailianghuang/FM-summary). Under the assumption that (a) there is one causal variant in each locus, and (b) the causal variant is observed in the genotype data, the credible set can be considered to have a 99% probability of containing the causal variant. For each the 12 genome-wide significant loci, variants within 1MB and in LD with correlation r^2^ > 0.4 to the index variant were considered for inclusion in the credible set analysis. The credible set analysis was done using the European GWAS meta-analysis to ensure consistent LD structure in the analyzed cohorts.

### Biological annotation of variants in credible set

The variants in the credible set for each locus, were annotated based on external reference data in order to evaluate potential functional consequences. In particular, we identify: (a) Gene and regulatory consequences annotated by Variant Effect Predictor (VEP) using Ensembl with genome build GRCh37^108^. We exclude upstream and downstream consequences, and consequences for transcripts that lack a HGNC gene symbol (e.g. vega genes). (b) Variants within 2kb upstream of the transcription start site (TSS) of at least one gene isoform based on Gencode v19^109^. (c) Variants annotated as interacting with a given gene in Hi-C data from samples of developing human cerebral cortex during neurogenesis and migration^110^. Annotations are considered for both the germinal zone (GZ), primarily consisting of actively dividing neural progenitors, and the cortical and subcortical plate (CP), primarily consisting of post-mitotic neurons. (d) Variants identified as eQTLs based on gene expression in GTEx^111^ or BIOS^73^. Expression quantitative trait loci were annotated using FUMA (http://fuma.ctglab.nl/). We restricted to eQTL associations with false discovery fate (FDR) < 1e-3 within each dataset. (e) Chromatin states of each variant based on the 15-state chromHMM analysis of epigenomics data from Roadmap^112^. The 15 states are summarized to annotations of active chromatin marks (i.e. Active TSS, Flanking Active TSS, Flanking Transcription, Strong Transcription, Weak Transcription, Genic Enhancer, Enhancer, or Zinc Finger [ZNF] gene), chromatin marks (Heterochromatin, Bivalent TSS, Flanking Bivalent TSS, Bivalent Enhancer, Repressed Polycomb, or Weak Repressed Polycomb), or quiescent. The most common chromatin state across 127 tissue/cell types was annotated using FUMA (http://fuma.ctglab.nl/). We also evalauted the annotated chromatin state from fetal brain.

### Gene-set analyses

Gene-based association with ADHD was estimated with MAGMA 1.05^83^ using the summary statistics from the European GWAS meta-analysis (N_cases_ = 19,099; N_controls_ = 34,194; See Supplementary Information, Supplementary Information Table 1). Association was tested using the SNP-wise mean model, in which the sum of -log(SNP P-value) for SNPs located within the transcribed region (defined using NCBI 37.3 gene definitions) was used as test statistic. MAGMA accounts for gene-size, number of SNPs in a gene and LD between markers when estimating gene-based P-values. LD correction was based on estimates from the 1000 genome phase 3 European ancestry samples^34^.

The generated gene-based P-values were used to analyze sets of genes in order to test for enrichment of association signals in genes belonging to specific biological pathways or processes. In the analysis only genes on autosomes, and genes located outside the broad MHC region (hg19:chr6:25-35M) were included. We used the gene names and locations and the European genotype reference panel provided with MAGMA. For gene sets we used sets with 10-1000 genes from the Gene Ontology sets^81^ currated from MsigDB 6.0^82^.

Targeted *FOXP2* downstream target gene sets were analysed for association with ADHD. Three sets were examined: 1) Putative target genes of *foxp2* that were enriched in wild type compared to control *foxp2* knockout mouse brains in ChIP-chip experiments (219 genes), 2) Genes showing differential expression in wild type compared to *foxp2* knockout mouse brains (243 genes), and 3) *FOXP2* target genes that were enriched in either or both basal ganglia (BG) and inferior frontal cortex (IFC) from human fetal brain samples in ChIP-chip experiments (258 genes). Curated short lists of high-confidence genes were obtained from Vernes et al.^84^ and Spiteri et al^85^.

A set of evolutionarily highly constrained genes were also analysed. The set of highly constrained genes was defined using a posterior probability of being loss-of-function intolerant (pLI) based on the observed and expected counts of protein-truncating variants (PTV) within each gene in a large study of over 60,000 exomes (the Exome Aggregation Consortium; ExAC)^86^. Genes with pLI ≥0.9 were selected as the set of highly constrained genes (2932 genes).

### Replication of GWAS loci

To replicate the results of the ADHD GWAS meta-analysis we compared the results to analyses from 23andMe and EAGLE. We evaluated evidence for replication based on: (a) genetic correlation between the ADHD GWAS and each replication cohort; (b) sign tests of concordance between the ADHD GWAS meta-analysis and each replication cohort; (c) meta-analysis of the ADHD GWAS meta-analysis results with the results from analyses of the replication cohorts; and (d) tests of heterogeneity in the meta-analyses of the ADHD GWAS meta-analysis with the replication cohorts.

Genetic correlations were calculated using LD score regression^37^ with the same procedure as described above. For the sign test, LD clumping was performed for all variants with P < 1 x 10^−4^ in the ADHD GWAS meta-analysis using LD estimated from European ancestry individuals from 1000 Genomes Phase 3 data. The proportion of variants with a concordant direction of effect in the two replication samples (*π*) was evaluated using a one sample test of the proportion with Yates’ continuity correction against a null hypothesis of *π* = 0.50 (i.e. the signs are concordant between the two analyses by chance). This test was done for loci passing P-value thresholds of P < 5 x 10^−8^, P < 1 x 10^−7^, P < 1x 10^−6^, P < 1 x 10^−5^, and P < 1 x 10^−4^ in the ADHD GWAS meta-analysis (see Supplementary Information).

We performed three meta-analyses based on the ADHD GWAS meta-analysis result and the results from the two replication cohorts. First, we performed an inverse variance-weighted meta-analysis of the ADHD GWAS meta-analysis with the results of the 23andMe GWAS of self-reported ADHD case status. Second, we performed a meta-analysis combining the results from clinically ascertained ADHD with results from GWAS of ADHD-related behavior in childhood population samples (the EAGLE data). This was done using a modified sample size-based weighting method (see below). Third, we applied the modified sample size-based weighting method to meta-analyze the EAGLE GWAS with the ADHD+23andMe GWAS meta-analysis.

For meta-analyses including the EAGLE cohort, modified sample size-based weights were derived to accounts for the respective heritabilities, genetic correlation, and measurement scale of the GWASs (Supplementary Information). To summarize, given *z*-scores *Z*_*1j*_ and *Z*_*2j*_ resulting from GWAS of SNP *j* in a dichotomous phenotype (e.g. ADHD) with sample size *N*_*I*_ and a continuous phenotype (e.g. ADHD-related traits) with sample size *N*_*2*_, respectively, we calculate

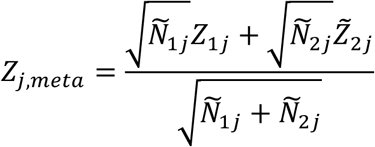

where

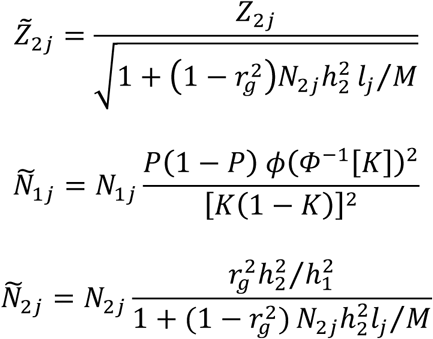

The adjusted sample sizes 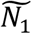 and 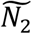 reflect differences in power between the studies due to measurement scale and relative heritability that is not captured by sample size. The calculation of 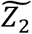 reduces the contribution of the continuous phenotype’s GWAS to the meta-analysis based on imperfect genetic correlation with the dichotomous phenotype of interest (i.e. ADHD). The adjustments are computed based on the sample prevalence (*P*) and population prevalence (*K*) of the dichotomous phenotype, the estimated liability scale SNP heritability of the two phenotypes (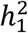 and 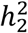), and the genetic correlation (*r*_*g*_) between the two phenotypes, as well as the average SNP LD score (*l*_*j*_) and the number of SNPs (*M*). Heritability and genetic correlation values to compute these weights are computed using LD score regression. This meta-analysis weighting scheme is consistent with weights alternatively derived based on modelling the joint distribution of marginal GWAS beta across traits^113^.

To test heterogeneity with each replication cohort, we considered Cochran’s *Q* test of heterogeneity in the first two replication meta-analyses described above. Specifically, we evaluated the one degree of freedom test for heterogeneity between the ADHD GWAS meta-analysis and the replication cohort.

## Acknowledgements

The iPSYCH team acknowledges funding from the Lundbeck Foundation (grant no R102-A9118 and R155-2014-1724), the Stanley Medical Research Institute, the European Research Council (project no: 294838), the European Community’s Horizon 2020 Programme (H2020/2014-2020) under Grant No. 667302 (CoCA), the Novo Nordisk Foundation for supporting the Danish National Biobank resource, and grants from Aarhus and Copenhagen Universities and University Hospitals, including support to the iSEQ Center, the GenomeDK HPC facility, and the CIRRAU Center.

The Broad Institute and Massachusetts General Hospital investigators would like to acknowledge support from the Stanley Medical Research Institute and NIH grants: 5U01MH094432-04(PI: Daly), 1R01MH094469 (PI: Neale), 1R01MH107649-01 (PI: Neale), 1R01MH109539-01 (PI: Daly). We thank T., Lehner, A. Addington and G. Senthil for their support in the Psychiatric Genomics Consortium. Statistical analyses were carried out on the Genetic Cluster Computer (http://www.geneticcluster.org) hosted by SURFsara and financially supported by the Netherlands Scientific Organization (NWO 480-05-003) along with a supplement from the Dutch Brain Foundation and the VU University Amsterdam. The GRAS data collection was supported by the Max Planck Society, the Max-Planck-Förderstiftung, and the DFG Center for Nanoscale Microscopy & Molecular Physiology of the Brain (CNMPB), Göttingen, Germany.

Dr. J. Martin was supported by the Wellcome Trust (Grant No: 106047).

Dr. Faraone is supported by the K.G. Jebsen Centre for Research on Neuropsychiatric Disorders, University of Bergen, Bergen, Norway, the European Union’s Seventh Framework Programme for research, technological development and demonstration under grant agreement no 602805, the European Union’s Horizon 2020 research and innovation programme under grant agreement No 667302 and NIMH grants 5R01MH101519 and U01 MH109536-01.

The Yale-Penn site study was supported by National Institutes of Health Grants RC2 DA028909, R01 DA12690, R01 DA12849, R01 DA18432, R01 AA11330, and R01 AA017535 and the Veterans Affairs Connecticut and Philadelphia Veterans Affairs Mental Illness Research, Educational, and Clinical Centers. Genotyping services for a part of our genome-wide association study were provided by the Center for Inherited Disease Research and Yale University (Center for Genome Analysis). Center for Inherited Disease Research is fully funded through a Federal contract from the National Institutes of Health to The Johns Hopkins University (contract number N01-HG-65403).

Dr. Haavik is supported by grants from Stiftelsen K.G. Jebsen, University of Bergen and The Research Council of Norway.

Dr. Cormand received financial support for this research from the Spanish ‘Ministerio de Economía y Competitividad’ (SAF2015-68341-R) and ‘Generalitat de Catalunya/AGAUR’ (2014SGR932). Dr. Cormand, Dr. Reif and collaborators received funding from the European Community’s Seventh Framework Programme (under grant agreement number 602805, Aggressotype), the European Community’s H2020 Programme (under grant agreements number 667302, CoCA, and 402003, MiND), the ECNP network ‘ADHD across the lifespan’ and DFG CRC 1193, subproject Z03. Dr. Andreassen is supported by the Research Council of Norway (grant nos: 223273, 248778, 213694, 249711), and KG Jebsen Stiftelsen.

Dr. Kuntsi’s research on ADHD is supported by the European Commission (grant agreements no. 643051 MiND, 667302 CoCA and 602805 Aggressotype); Action Medical Research (GN2080 and GN2315); 4 Medical Research Council and SGDP Centre PhD studentships; and by the ECNP Network ADHD Across the Lifespan. Dr. Langley was funded by Wellcome Trust (Grant No: 079711)

Dr. Thapar received ADHD funding from the Wellcome Trust, Medical Research Council (MRC UK), Action Medical Research.

Barbara Franke’s research is supported by funding from a personal Vici grant of the Netherlands Organisation for Scientific Research (NWO; grant 016-130-669, to BF), from the European Community’s Seventh Framework Programme (FP7/2007 – 2013) under grant agreements n° 602805 (Aggressotype), n° 602450 (IMAGEMEND), and n° 278948 (TACTICS), and from the European Community’s Horizon 2020 Programme (H2020/2014 – 2020) under grant agreements n° 643051 (MiND) and n° 667302 (CoCA). In addition, this work was supported by the European College of Neuropsychopharmacology (ECNP Network “ADHD across the Lifespan”).

Dr. Schachar received support from Bank Chair in Child Psychiatry, Canadian Institutes of Health Research (MOP-106573 and MOP – 93696).

Dr. Roussos was supported by the National Institutes of Health (R01AG050986 Roussos and R01MH109677), Brain Behavior Research Foundation (20540), Alzheimer’s Association (NIRG-340998) and the Veterans Affairs (Merit grant BX002395).

We thank the customers of 23andMe who answered surveys, as well as the employees of 23andMe, who together made this research possible.

We gratefully acknowledge all the studies and databases that made GWAS summary data available: ADIPOGen (Adiponectin genetics consortium), C4D (Coronary Artery Disease Genetics Consortium), CARDIoGRAM (Coronary ARtery DIsease Genome wide Replication and Meta-analysis), CKDGen (Chronic Kidney Disease Genetics consortium), dbGAP (database of Genotypes and Phenotypes), DIAGRAM (DIAbetes Genetics Replication And Meta-analysis), ENIGMA (Enhancing Neuro Imaging Genetics through Meta Analysis), EAGLE (EArly Genetics & Lifecourse Epidemiology Consortium, excluding 23andMe), EGG (Early Growth Genetics Consortium), GABRIEL (A Multidisciplinary Study to Identify the Genetic and Environmental Causes of Asthma in the European Community), GCAN (Genetic Consortium for Anorexia Nervosa), GEFOS (GEnetic Factors for OSteoporosis Consortium), GIANT (Genetic Investigation of ANthropometric Traits), GIS (Genetics of Iron Status consortium), GLGC (Global Lipids Genetics Consortium), GPC (Genetics of Personality Consortium), GUGC (Global Urate and Gout consortium), HaemGen (haemotological and platelet traits genetics consortium), HRgene (Heart Rate consortium), IIBDGC (International Inflammatory Bowel Disease Genetics Consortium), ILCCO (International Lung Cancer Consortium), IMSGC (International Multiple Sclerosis Genetic Consortium), MAGIC (Meta-Analyses of Glucose and Insulin-related traits Consortium), MESA (MultiEthnic Study of Atherosclerosis), PGC (Psychiatric Genomics Consortium), Project MinE consortium, ReproGen (Reproductive Genetics Consortium), SSGAC (Social Science Genetics Association Consortium) and TAG (Tobacco and Genetics Consortium), TRICL (Transdisciplinary Research in Cancer of the Lung consortium), UK Biobank.

We gratefully acknowledge the contributions of Alkes Price (the systemic lupus erythematosus GWAS and primary biliary cirrhosis GWAS) and Johannes Kettunen (lipids metabolites GWAS).

## Author Contributions

This section will be filled in the review process

## Disclosures

In the past year, Dr. Faraone received income, potential income, travel expenses continuing education support and/or research support from Lundbeck, Rhodes, Arbor, KenPharm, Ironshore, Shire, Akili Interactive Labs, CogCubed, Alcobra, VAYA, Sunovion, Genomind and Neurolifesciences. With his institution, he has US patent US20130217707 A1 for the use of sodium-hydrogen exchange inhibitors in the treatment of ADHD. In previous years, he received support from: Shire, Neurovance, Alcobra, Otsuka, McNeil, Janssen, Novartis, Pfizer and Eli Lilly. Dr. Faraone receives royalties from books published by Guilford Press: Straight Talk about Your Child’s Mental Health, Oxford University Press: Schizophrenia: The Facts and Elsevier: ADHD: Non-Pharmacologic Interventions. He is principal investigator of www.adhdinadults.com.

Dr. Neale is a member of Deep Genomics Scientific Advisory Board and has received travel expenses from Illumina. He also serves as a consultant for Avanir and Trigeminal solutions.

Dr. Rohde has received honoraria, has been on the speakers’ bureau/advisory board and/or has acted as a consultant for Eli-Lilly, Janssen-Cilag, Novartis, Medice and Shire in the last three years. He receives authorship royalties from Oxford Press and ArtMed. He also received travel award for taking part of 2015 WFADHD meeting from Shire. The ADHD and Juvenile Bipolar Disorder Outpatient Programs chaired by him received unrestricted educational and research support from the following pharmaceutical companies in the last three years: Eli-Lilly, Janssen-Cilag, Novartis, and Shire.

Over the last three years Dr. Sonuga-Barke has received speaker fees, consultancy, research funding and conference support from Shire Pharma and speaker fees from Janssen Cilag. He has received consultancy fees from Neurotech solutions, Aarhus University, Copenhagen University and Berhanderling, Skolerne, Copenhagen, KU Leuven. Book royalties from OUP and Jessica Kingsley. He is the editor-in-chief of the Journal of Child Psychology and Psychiatry for which his University receives financial support.

Barbara Franke has received educational speaking fees from Merz and Shire.

Dr. Schachar’s disclosures: ehave equity and advisory board, Ironshore Pharmaceuticals Advisory Board.

Dr. Reif has received a research grant from Medice, and speaker’s honorarium from Medice and Servier.

**Correspondence and requests for materials should be addressed to:**

Benjamin Neale

Anders D. Børglum

Stephen Faraone

## Extended Data

**eTable 1.** Bayesian credible sets of variants for each of the 12 genome-wide significant loci

**eTable 2.** Summary of the observed annotations for the credible set at each genome-wide significant locus

**eTable 3.** Variant-level annotations for the credible set at each genome-wied significant locus

**eTable 4.** Results of gene set analyses using sets from Gene Ontology

**eTable 5.** Extended results from genetic correlation analyses of ADHD and 220 phenotypes

**eTable 6.** Genome-wide significant index variants in meta-analyses of iPSYCH, PGC, 23andMe and EAGLE

